# Reduced H^+^ channel activity disrupts pH homeostasis and calcification in coccolithophores at low ocean pH

**DOI:** 10.1101/2021.09.06.458185

**Authors:** Dorothee Kottmeier, Abdul Chrachri, Gerald Langer, Katherine Helliwell, Glen L. Wheeler, Colin Brownlee

## Abstract

Coccolithophores produce the bulk of ocean biogenic calcium carbonate but this process is predicted to be negatively affected by future ocean acidification scenarios. Since coccolithophores calcify intracellularly, the mechanisms through which changes in seawater carbonate chemistry affect calcification remain unclear. Here we show that voltage-gated H^+^ channels in the plasma membrane of *Coccolithus braarudii* serve to regulate pH and maintain calcification under normal conditions, but have greatly reduced activity in cells acclimated to low pH. This disrupts intracellular pH homeostasis and impairs the ability of *C. braarudii* to remove H^+^ generated by the calcification process, leading to specific coccolith malformations. These coccolith malformations can be reproduced by pharmacological inhibition of H^+^ channels. Heavily-calcified coccolithophore species such as *C. braarudii*, which make the major contribution to carbonate export to the deep ocean, have a large intracellular H^+^ load and are likely to be most vulnerable to future decreases in ocean pH.

## Introduction

Anthropogenic CO_2_ emissions and the subsequent dissolution of CO_2_ in seawater have resulted in substantial changes in ocean carbonate chemistry ^1^. The resultant decrease in seawater pH, termed ocean acidification, is predicted to be particularly detrimental for calcifying organisms ^2^. Mean global surface ocean pH is predicted to fall as low as 7.7 by 2100 ^3^ and is likely to continue to fall further in the following centuries. Present day marine organisms can experience significant fluctuations in seawater pH, particularly in coastal and upwelling regions ^4, 5^. Ocean acidification is therefore predicted to have an important influence not only on mean surface ocean pH, but also on the extremes of pH experienced by marine organisms ^6, 7^. Coccolithophores, characterised by their covering of intricately-formed calcite scales (coccoliths), account for the bulk of global biological calcification and around 20% of ocean productivity, making major contributions to global biogeochemical cycles, including the long-term export of both inorganic and organic carbon from the ocean photic zone to deep waters ^8, 9^. Unlike the vast majority of calcifying organisms, coccolithophore calcification occurs in an intracellular compartment, the Golgi-derived coccolith vesicle (CV), effectively isolating the calcification process from direct changes in seawater carbonate chemistry. Nevertheless, extensive laboratory observations indicate that ocean acidification may negatively impact coccolithophore calcification, albeit with significant variability of responses between species and strains ^10-14^. The negative impact on calcification rates occurs at calcite saturation states (Ω) >1, indicating it results primarily from impaired cellular production rather than dissolution ^10, 15^. However, prediction of how natural coccolithophore populations may respond to future changes in ocean pH are hampered by lack of mechanistic understanding of pH impacts at the cellular level ^10^.

As calcification occurs intracellularly using external HCO_3_^-^ as the primary dissolved inorganic carbon (DIC) source ^16-18^, coccolith formation is not directly dependent on external CO_3_^2-^ concentrations. However, the uptake of HCO_3_^-^ as a substrate for calcification results in the equimolar production of CaCO_3_ and H^+^ in the CV ^18^. In order to maintain saturation conditions for calcite formation, H^+^ produced by the calcification process must be rapidly removed from the CV, placing extraordinary demands for cellular pH regulation to prevent cellular acidosis ^18^.

Lower calcification rates under ocean acidification conditions appear to be primarily due to decreased pH rather than other aspects of carbonate chemistry ^10, 19, 20^. Coccolithophores exhibit highly unusual membrane physiology, including the presence of voltage-gated H^+^ channels in the plasma membrane ^21^ and a high sensitivity of cytosolic pH (pH_cyt_) to changes in external pH (pH_o_) ^21, 22^. Voltage-gated H^+^ channels are associated with rapid H^+^ efflux in a number of specialised animal cell types ^23^ and contribute to effective pH regulation in coccolithophores ^21^. As H^+^ channel function is dependent on the electrochemical gradient of H^+^ across the plasma membrane, this mechanism could be impaired under lower seawater pH. However, it remains unknown whether H^+^ channels play a direct role in removal of calcification-derived H^+^ or contribute to the sensitivity of coccolithophores to ocean acidification.

Coccolithophores exhibit significant diversity in their extent of calcification (Supplementary Fig. 1). The abundant bloom forming species *Emiliania huxleyi* is moderately calcified (ratio of particulate inorganic carbon to particulate organic carbon (PIC/POC) of around 1) and has been the focus of the vast majority of the studies into the effects of environmental change in coccolithophores ^13^. Coastal species belonging to the Pleurochrysidaceae and Hymenomonadaceae are lightly calcified, commonly exhibiting a PIC/POC of less than 0.5 ^24-27^. Species such as *Coccolithus braarudii, Calcidiscus leptoporus* and *Helicosphaera carteri* exhibit much higher PIC/POC ratios and contribute the majority of carbonate export to the deep ocean in many areas ^28-30^. The physiological response of heavily-calcified coccolithophores to ocean acidification is therefore of considerable biogeochemical significance. Growth and calcification rates in *C. leptoporus* and *C. braarudii* are sensitive to pH values predicted to prevail on a future decadal timescale ^10, 15, 31, 32^. However, a mechanistic understanding of the different sensitivity of coccolithophore species to changing ocean carbonate chemistry is lacking.

The net H^+^ load in a cell is determined by the combination of metabolic processes that consume or produce H^+^. H^+^ fluxes in coccolithophores will be primarily determined by the balance of H^+^ consumed by photosynthesis and H^+^ generated by calcification, with uptake of different carbon sources a particularly important consideration (Fig. 1a). CO_2_ uptake for photosynthesis results in no net production or consumption of H^+^, whereas uptake of HCO_3_^-^ requires the equimolar consumption of H^+^ in order to generate CO_2_. Growth at elevated CO_2_ causes a switch from HCO_3_^-^ uptake to predominately CO_2_ uptake in *E. huxleyi* ^33, 34^. The associated net decrease in H^+^ consumption will therefore increase the H^+^ load in coccolithophores grown at elevated CO_2_, which may exacerbate the potential for cytosolic acidosis caused by lower seawater pH.

**Fig. 1.**
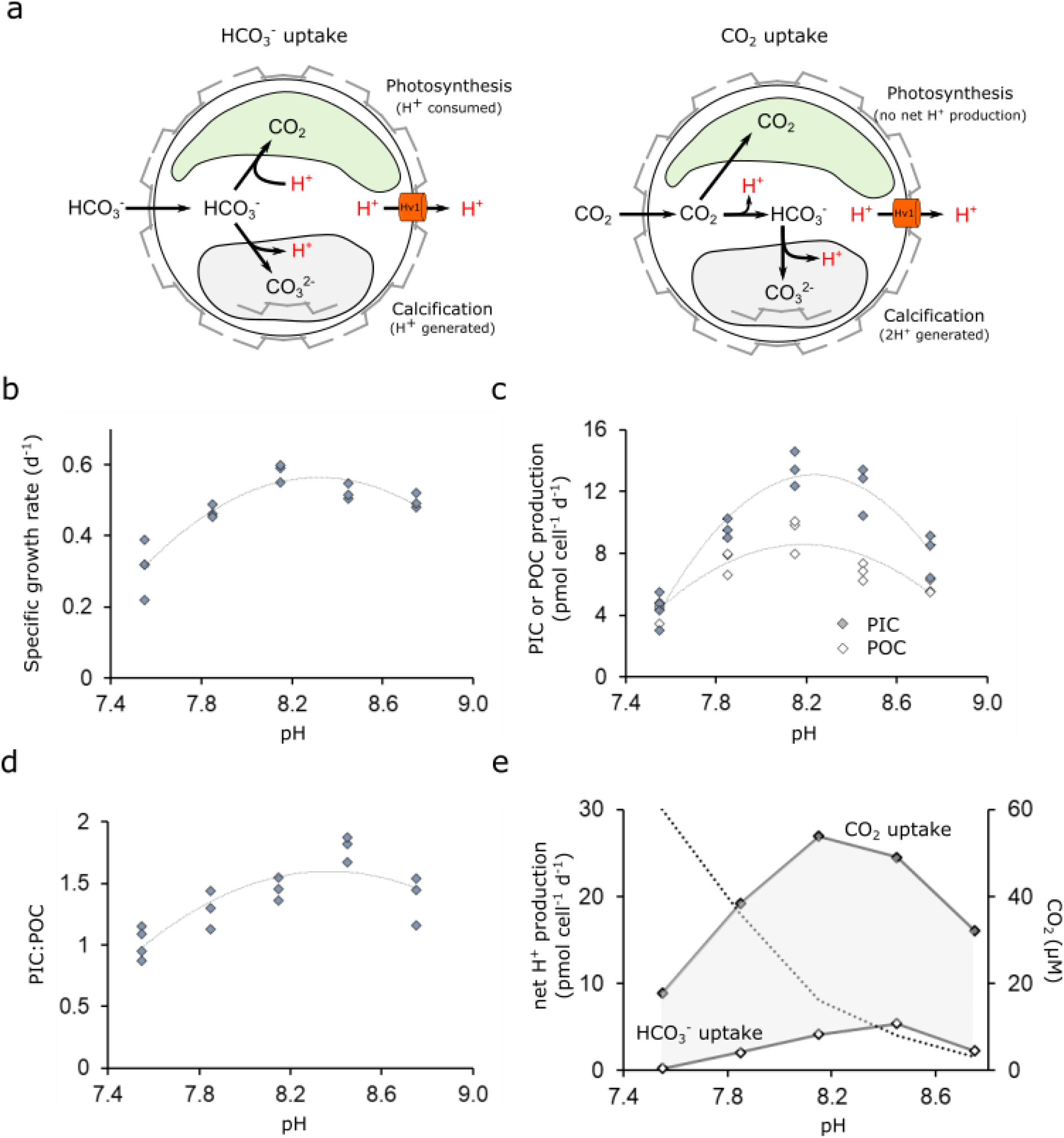
Physiology and H^+^ fluxes of *Coccolithus braarudii* cells grown at different seawater pH. **a** Schematic indicating H^+^ fluxes associated with photosynthesis and calcification in a coccolithophore cell. Whilst many metabolic processes may contribute to the cellular H^+^ budget, these two processes are likely to be the major contributors. In a cell taking up HCO_3_^-^, the H^+^ budget is balanced between H^+^ consumed during photosynthesis and H^+^ generated during calcification. In a cell taking up CO_2_, 2 H^+^ are produced for each molecule of CaCO_3_ produced and H^+^ are no longer consumed during photosynthesis. Excess H^+^ may be removed from the cell by H^+^ transporters in the plasma membrane, such as voltage-gated H^+^ channels (Hv). Coccolithophores take up both HCO_3_^-^ and CO_2_ across the plasma membrane, with increasing proportions of DIC taken up as CO_2_ as seawater CO_2_ increases ^34^. **b** Growth rate of *C. braarudii* cells acclimated to different seawater pH. n=3 replicates per treatment, line represents polynomial fit to mean. **c** Cellular production of particulate organic carbon (POC) through photosynthesis and particulate inorganic carbon (PIC) through calcification. The optima for both processes are close to the control conditions (pH 8.15). **d** As a consequence of the unequal changes in cellular POC and PIC production across the applied pH values, cellular PIC:POC ratios are minimal at pH 7.55 (≈1.0) and maximal at pH 8.45 (≈1.8). **e** Calculated net H^+^ budgets under the different pH regimes, based on rates of photosynthesis and calcification shown in (c) (See Methods). The concentration of CO_2_ in seawater is also shown (dashed line). Estimates are shown for cells using taking up only HCO_3_^-^ or only CO_2_. As *C. braarudii* cells will likely take up a mixture of both DIC species, with a shift towards greater CO_2_ usage at elevated CO_2_, the shaded area represents the potential range of H^+^ production. Regardless of DIC species used *C. braarudii* produces excess H^+^ at all applied pH values, but H^+^ production is much lower at pH 7.55 due to the decrease in calcification.

In this study we set out to better understand the cellular mechanisms underlying the sensitivity of coccolithophore calcification to lower pH. We subjected the heavily-calcified species *C. braarudii*, which is commonly found in temperate upwelling regions ^35, 36^, to conditions that reflect the range of pH values it may experience in current and future oceans. We show that acclimation to low pH leads to loss of H^+^ channel function and disruption of cellular pH regulation in *C. braarudii*. These effects are coincident with very specific defects in coccolith morphology that can be reproduced by direct inhibition of H^+^ channels. We conclude that H^+^ efflux through H^+^ channels is essential for maintaining both calcification rate and coccolith morphology. By providing a mechanistic insight into pH regulation during the calcification process, our results indicate that disruption of coccolithophore calcification in a future acidified ocean is likely to be most severe in heavily calcified species.

## Results

### Cellular H^+^ load varies with DIC use for calcification and photosynthesis

To examine more closely how the balance of photosynthesis and calcification may influence the cellular H^+^ load, we measured physiological parameters in *C. braarudii* cells acclimated to a broad range of carbonate chemistries (Supplementary Table 1). *C. braarudii* exhibited pH optima for growth, PIC and POC production of 8.32 ± 0.01, 8.20 ± 0.03 and 8.24 ± 0.02 respectively (pH_NBS_, n=3, ±SE), with sharp declines in these parameters exhibited by cells grown at pH 7.85 and 7.55 (Fig. 1b-d). PIC production decreased more strongly than POC production in acidified conditions, leading to lower PIC:POC ratios. These trends are in close agreement with other laboratory studies examining the response of *C. braarudii* to changing carbonate chemistries ^10, 15, 31, 32, 37^. Calculation of the H^+^ load from values of PIC and POC production indicated that H^+^ production by calcification exceeded H^+^ consumption by photosynthesis under all scenarios, being highest at optimal PIC:POC ratios (Fig. 1e). Although the large decrease in calcification rates at seawater pH 7.55 result in lower H^+^ production, the net H^+^ load could still be substantial due to a likely increase in CO_2_ uptake under these conditions (Fig. 1e) ^34^. The results illustrate that changes in the relative rates of photosynthesis and calcification, as well as in the carbon source used for photosynthesis, will have a major impact on the cellular H^+^ budget in *C. braarudii*, although in all cases there is a resultant requirement for net H^+^ efflux.

### Growth at low pH results in specific defects in coccolith morphology

Morphological defects in coccoliths are widely reported in coccolithophores grown under simulated ocean acidification conditions ^37, 38^, although there is little information on the specific nature of these malformations to aid mechanistic understanding of the impacts of low seawater pH on the calcification process. Scanning electron microscopy (SEM) analysis of coccolith morphology revealed that only 19.0 ± 5.0 % and 30.1 ± 2.7 % of coccoliths exhibited normal morphology at pH 7.55 and pH 7.85 respectively (n=3, ±SE) (Fig. 2a,b). Moreover, by performing a detailed categorisation of each morphological defect, we found a very distinct ‘type-pH’ malformation at low pH, in which the shield elements are malformed and greatly reduced in length so that the coccolith appears as a ring of calcite rather than a fully formed shield (Fig. 2b). This differs from an immature coccolith, in which the individual elements are reduced in length but correctly formed (Supplementary Fig. 2). Cells grown at low pH exhibited a large increase in the number of collapsed coccospheres observed by SEM analysis (Fig. 2c-d), indicating that the extensive malformations result in an inability to maintain the structural integrity of the coccosphere. Although the type-pH malformation has not been explicitly described previously, it can clearly be observed in other studies where *C. braarudii* has been grown at low pH ^37^. Importantly, type-pH malformations are distinct from coccolith malformations induced by other stressors, such as phosphate limitation or the Si analogue Ge ^39, 40^.

**Fig. 2.**
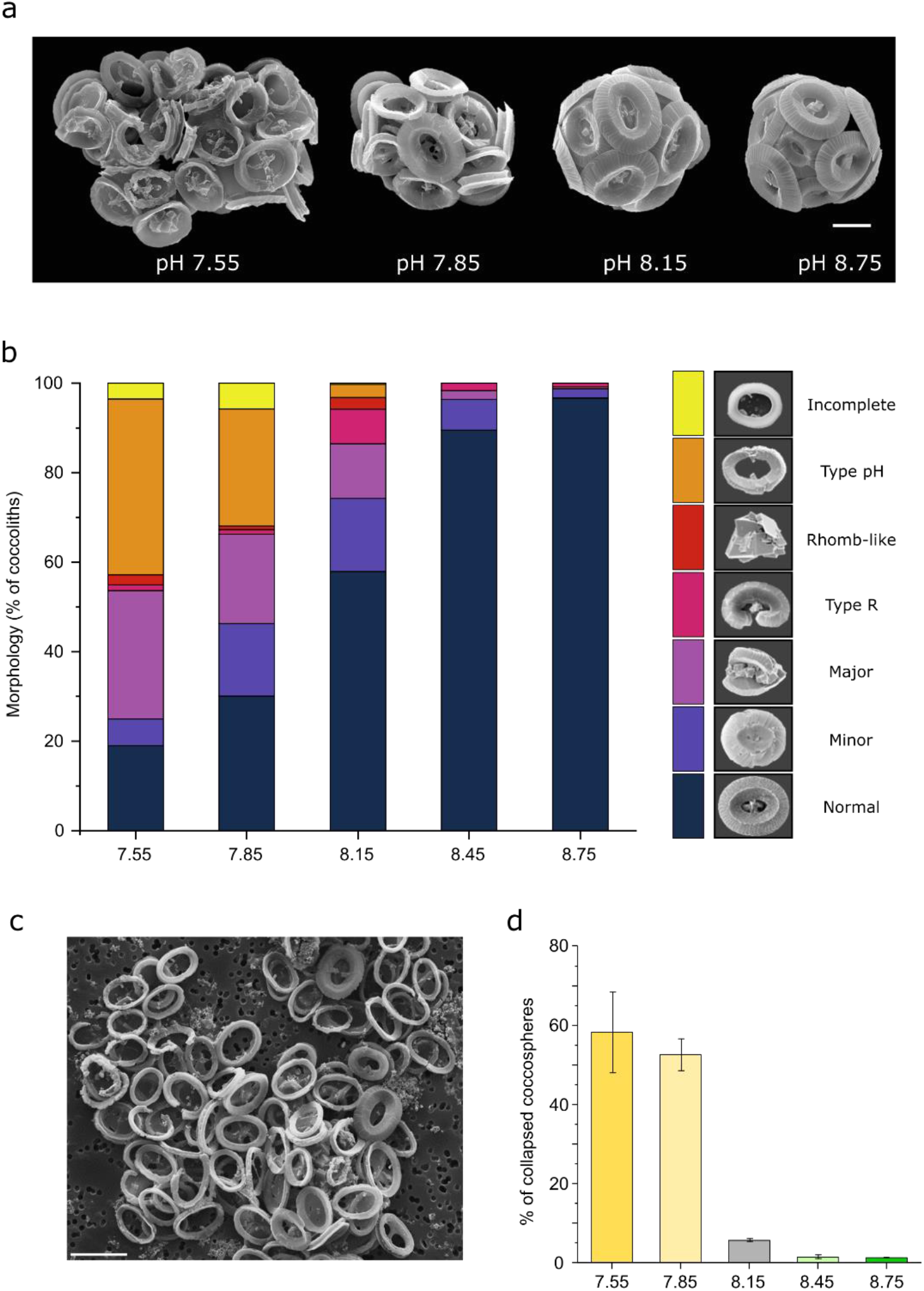
A unique defect in coccolith morphology occurs at low seawater pH. **A** Representative scanning electron micrographs of cells acclimated to pH 7.55, 7.85, 8.15 and 8.75. The majority of cells grown at pH ≥8.15 had intact coccospheres without crystal or coccolith malformation. The majority of cells grown under acidified conditions produced malformed coccoliths, resulting in abnormal or collapsed coccospheres. Bar = 5 μm **b** Quantification of coccolith malformations reveal an increasing proportion of defective coccoliths with seawater acidification. Coccoliths were grouped into morphological categories (see Methods). The morphological categories representing rhomb-like, R-type, major and minor malformations are commonly observed *C. braarudii* cells grown under various stressors ^73^. However, the distinct ‘type-pH’ was only observed in this study and appears unique to low pH (high CO_2_) conditions. The counts represent the mean of three independent replicates. A minimum of 350 coccoliths were counted for each replicate. **c** An example of *C. braarudii* cells grown at pH 7.55 exhibiting a high proportion of the distinctive ‘type-pH’ malformations. As the shield elements are not properly formed, the coccoliths are unable to interlock in the normal manner, resulting in the collapse of the coccospheres during preparation for SEM imaging. Bar = 10 μm. **d** The proportion of collapsed coccospheres increases at lower seawater pH. n= 3 replicates per treatment. Error bars represent SE.

### H^+^ channel function is greatly reduced following acclimation to lower pH

To investigate how these defects in coccolith morphology could arise, we examined the physiology of *C. braarudii* cells grown at low pH. *C. braarudii* exhibits an unusual large outwardly rectifying H^+^ current at membrane potentials positive of the H^+^ equilibrium potential (E_H+_), due to the activity of voltage-gated H^+^ channels in the plasma membrane ^21^. Our previous studies showed that the H^+^ channel activation potential tracked E_H+_ cross the plasma membrane. Calculation of the proton motive force (pmf) across the *C. braarudii* plasma membrane (at a resting membrane potential of -46 mV ^21^) indicates that there is a small net outward pmf at a seawater pH of 8.15 (Fig 3a). A decrease in pH_cyt_ results in a shift in the activation potential of the H^+^ current to more negative membrane potential values and increases the outwardly-directed pmf. These combined effects result in net H^+^ efflux and allow restoration of resting pH_cyt_ (Supplementary Fig. 3). However, at pH_o_ 7.55 the activation potential shifts to more positive values and the calculated pmf is no longer outward, so channel mediated H^+^ efflux would only occur following depolarisation of the membrane potential and/or further reductions in pH_cyt_ (Supplementary Figure 3).

**Fig. 3:**
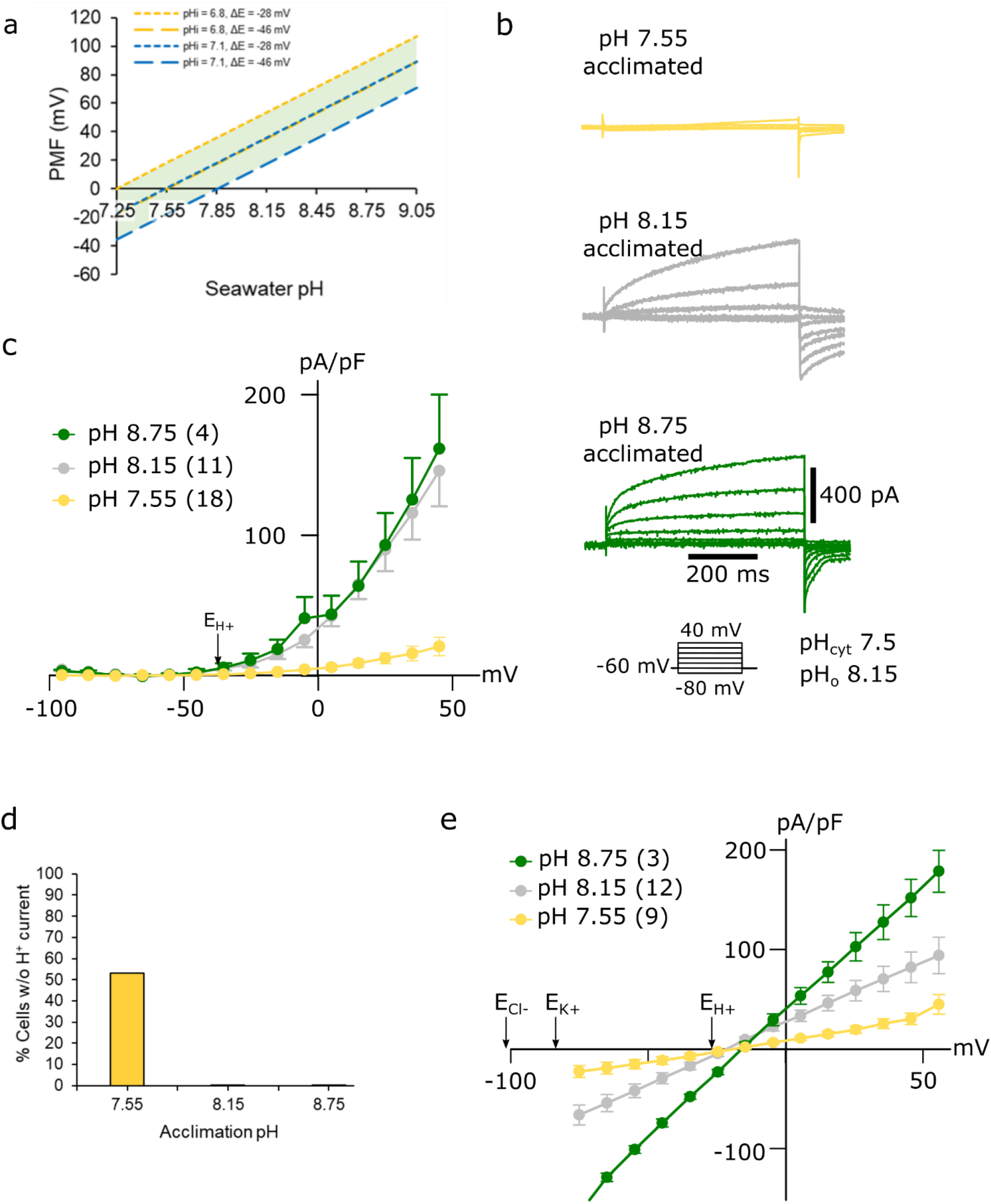
A reduced outward H^+^ current in *C. braarudii* cells acclimated to low seawater pH. **a** Estimation of the impact of changes in seawater pH on the proton motive force (PMF) across the plasma membrane. Models of PMF based on measured maximal or minimal pH_cyt_ (pH 6.8 and 7.1: See Fig. 4) in combination with measured minimal and maximal ΔE (−46mV and - 28mV, ^21^) suggest that PMF is close to zero at a seawater pH of approximately pH 7.55. Therefore passive H^+^ efflux via voltage-gated H^+^ channels becomes unfavourable, unless mediated by excursions of pH_cyt_ (lower cytosolic pH) or V_m_ (depolarisation). **b** Electrophysiological measurements of whole cell currents in response to incremental 1 s 10 mV depolarisations from -80 to +40 mV performed in artificial seawater buffered to pH 8.15. The large outward-directed ion current is predominately carried by H^+^. The maximal current is much smaller in cells acclimated to pH 7.55. Example of currents from a single cell are shown for each pH treatment. **c** Mean whole cell currents (plotted as pA/pF vs mV) for acclimated cells. Outward currents are observed when the plasma membrane is depolarised to potentials more positive than the equilibrium potential of H^+^ (E_H+_; arrow). Cells acclimated to pH 7.55 exhibit a greatly reduced outward current. Values in parentheses represent n, bars represent SE. **d** The proportion of cells that do not show an outward current (defined as an outward current <2.5 pA/pF at +45 mV) is greatly increased in cultures acclimated to a seawater pH of 7.55 compared to cultures acclimated to pH 8.15 or 8.75. n=18. **e** Tail current analysis indicating that reversal potentials (E_rev_) are close to E_H+_ and more positive than the E_K+_ and E_Cl-_ in all treatments, suggesting that the observed currents in all treatments are predominately carried by H^+^.

In order to determine the impact of growth at unfavourable seawater pH on H^+^ channel function, we monitored H^+^ currents using patch clamp recordings in C. *braarudii* cells previously acclimated to pH_o_ 7.55, 8.15 or 8.75. The mean amplitude of the outward H^+^ current, when measured at pH_o_ 8.15, was greatly reduced in cells that had been acclimated to pH 7.55 (Fig. 3b-d). 52.9% of cells acclimated to pH 7.55 exhibited either greatly reduced or undetectable outward current (Fig. 3d), although these cells still displayed inward Cl^-^ currents typical of healthy *C. braarudii* cells ^41^ (Supplementary Fig.4). The outward currents exhibited a reversal potential (E_rev_) close to E_H+_, indicating that H^+^ remained as the primary charge carrier in all cases (Fig. 3e). The results suggest that acclimation of *C. braarudii* to a low seawater pH unsuited to the operation of H^+^ channels results in the loss of H^+^ channel function.

### A unique family of H^+^ channels in calcifying coccolithophores

Homologues of the mammalian voltage-gated H^+^ channel, Hv1, are present in coccolithophores and a range of other phytoplankton, although the large outward H^+^ currents typical of *C. braarudii* have not been reported in other algal cells, suggesting that H^+^ channels are utilised for alternative roles in non-calcifying cells (e.g. in supporting NADPH oxidase activity ^42^ or in dinoflagellate bioluminescence ^43, 44^). We previously characterised Hv1 channels from *E. huxleyi* and *C. braarudii* ^21^. Further analysis of haptophyte transcriptomes ^45^ revealed that coccolithophores possess an additional H^+^ channel homologue (Hv2) that was not found in non-calcifying haptophytes (Supplementary Fig.5). *C. braarudii* Hv2 exhibited robust H^+^ currents when expressed in a heterologous system (Supplementary Fig.5). The presence of this additional homologue suggests that coccolithophore H^+^ channels have undergone functional specialisation related to calcification. In support of a specific role in calcification, we found that *HV1* and *HV2* were only expressed in the heavily calcified heterococcolith-bearing diploid life cycle phase of *C. braarudii* and were not detected in the lightly calcified holococcolith-bearing haploid life cycle phase (Supplementary Fig.6). However, we did not find any significant change in the expression of either *HV1* or *HV2* in diploid cells acclimated to low pH (Supplementary Fig.6). This suggests that the greatly reduced H^+^ conductance in these cells results from post-transcriptional or post-translational regulation of H^+^ channels.

### pH homeostasis is disrupted at low seawater pH

To determine whether the reduced H^+^ channel activity in cells acclimated to low pH led to disrupted cellular pH homeostasis, we examined resting pH_cyt_. Cells acclimated to pH 7.55 exhibited a significantly lower mean pH_cyt_ values than cells acclimated to pH 8.15 or pH 8.75 (Fig. 4a, Supplementary Fig.7). Cells acclimated to pH 8.15 or 8.75 retained the ability to adjust intracellular pH rapidly within seconds when exposed to a higher or lower pH ^21, 22^, but this response was greatly reduced in cells acclimated to pH 7.55 (Fig. 4b-d). To determine the capacity for H^+^ efflux, we transiently exposed cells to pH 6.5 and examined their ability to restore pH_cyt_ on transfer to pH 8.15. Nearly all cells acclimated to pH 8.15 or 8.75 showed a substantial decrease in cytosolic [H^+^] on transfer from pH 6.5 to 8.15 (Fig. 4e). However, many cells acclimated to pH 7.55 showed little or no capacity to lower cellular [H^+^] on transfer from pH 6.5 to 8.5, indicating the presence of distinct populations of responsive and unresponsive cells. (Fig. 4e-f). Thus, a significant proportion of cells acclimated to pH 7.55 exhibit a defect in H^+^ efflux, which likely reflects the highly reduced H^+^ channel activity in these cells (Fig. 3d).

**Fig. 4:**
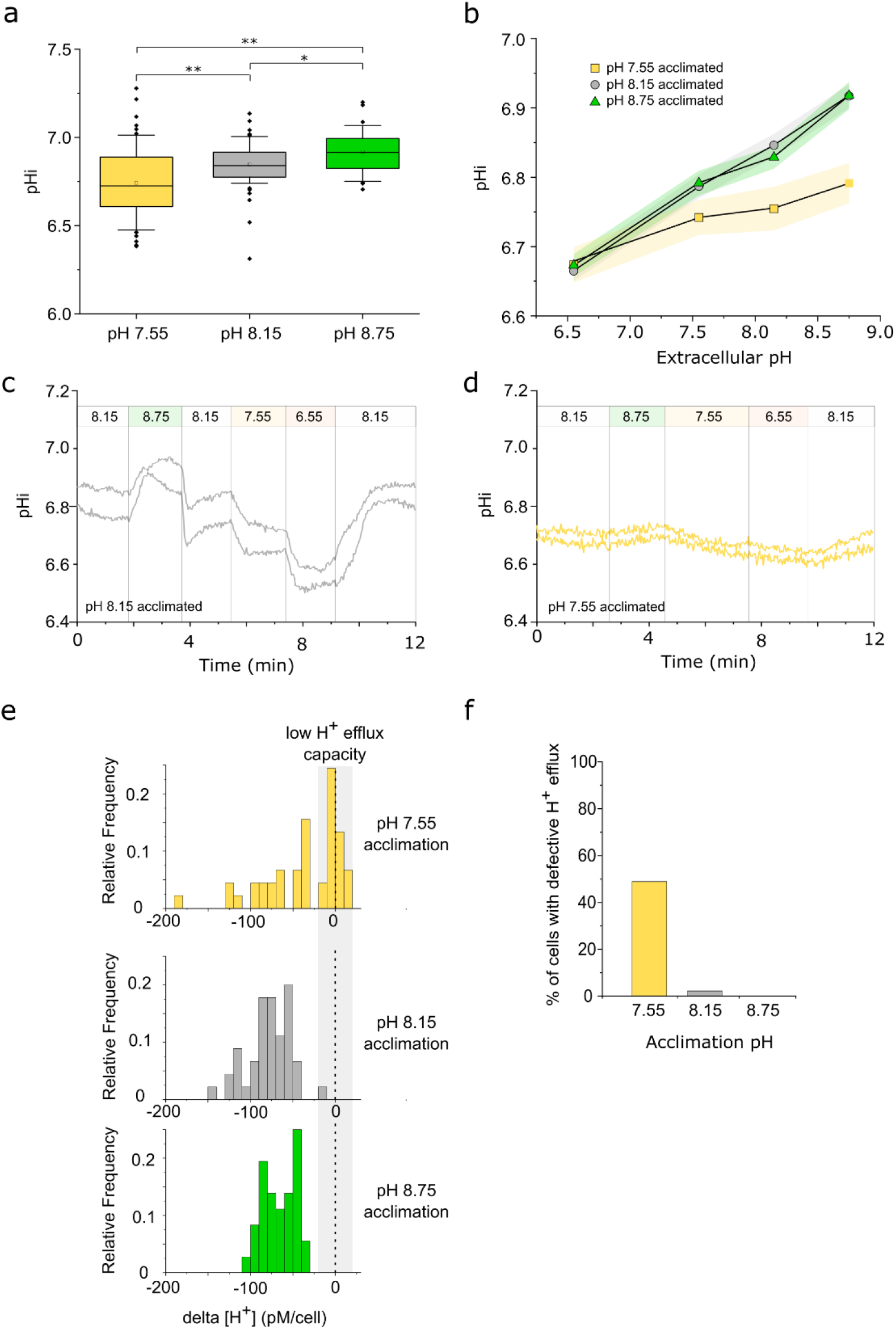
Changes in intracellular pH (pH_cyt_) in response to seawater acidification and alkalinisation. **a** Intracellular pH (pH_cyt_) measured at acclimation pH conditions in *C. braarudii* cells loaded with the pH-responsive fluorescent dye SNARF-AM. Cells acclimated to pH 7.55 show a lower mean pH_cyt_ to those acclimated to pH 8.15 and pH 8.75. n= 63, 61 and 36 cells for pH 7.55, 8.15 and 8.75 respectively. 1-way ANOVA, Holm-Sidak post-hoc test, * p<0.05, ** p<0.01. Box plots indicate mean (open square), median, 25-75^th^ percentiles (box) and 10-90^th^ percentiles (whiskers). **b** pH_cyt_ regulation following rapid changes in external pH. Acclimated cells were perfused with seawater at pH 6.55, 7.55, 8.15 and 8.75 for 2 minutes each to examine their ability to regulate pH_cyt_. Cells acclimated to pH 8.15 and 8.75 show the rapid adjustment of pH_cyt_ typical of coccolithophore cells ^21,22^. However, cells acclimated to pH 7.55 show a much lower change in pH_cyt_ (n=25, 61 and 36 respectively for pH 7.55, 8.15 and 8.75). Shaded areas represent SE. **c** Example of rapid changes in intracellular pH (pH_cyt_) in cells acclimated to pH 8.15. Cells were perfused with ASW pH 8.15 and pHi was monitored as the perfusion was switched to a higher or lower pH. Two representative cells are shown. **d** Example of cells acclimated to pH 7.55 exhibiting little change in pH_cyt_ following changes in external pH. Note that the time course of the perfusion differs slightly from that shown in (C). Two representative cells are shown. **e** Detailed examination of pH_cyt_ recovery during a transition from seawater pH 6.5 to seawater pH 8.15. The frequency histogram indicates the change in pH_cyt_ (shown as Δ[H^+^]) in individual cells acclimated to different pH regimes. Whilst nearly all cells acclimated to pH 8.15 and 8.75 exhibit a substantial decrease in [H^+^] on transfer from pH 6.5 to higher pH, many cells acclimated to pH 7.55 are unable to respond, indicative of a defect in H^+^ efflux. n=55, 61 and 36 cells). Cells acclimated to pH 7.55 exhibit a significantly different distribution to pH 8.15 or 8.75 (2-sample Kolmogorov-Smirnov test, p<0.05). **f** Proportions of cells exhibiting defective H^+^ efflux in the experiment described in (E). Defective H^+^ efflux was defined as a Δ[H^+^] less than 20 pM.

### Pharmacological inhibition of H^+^ channel function disrupts coccolith morphology

Our results suggest that loss of H^+^ channel function and subsequent disruption of pH homeostasis is directly responsible for the defects in calcification exhibited by *C. braarudii* grown at low pH. To directly test this hypothesis, we treated cells with two inhibitors of Hv channels, Zn^2+ 23^ and 2-guanidinobenzimidazole (2-GBI) ^46^. Cells grown in 35 μM Zn^2+^, which inhibits the outward H^+^ conductance in *C. braarudii* by approximately 60% ^21^, showed only a small reduction in growth rate (control 0.54 ± 0.01 d^-1^ compared to Zn^2+^-treated 0.47 ± 0.01 d^- 1^, n=3, se) (Fig. 5a). However, SEM examination of Zn^2+^-treated cells after 5 d revealed severe disruptions of coccolith morphology (Fig. 5b-d). Importantly, Zn-treated cells exhibited the unique type-pH coccolith malformations, which were completely absent from control cells. Growth of cells in 15 μM 2-GBI for 5 d also resulted in the presence of type-pH coccolith malformations (Supplementary Fig.8), suggesting that this calcification phenotype is specifically associated with impaired H^+^ channel function. Our results show that pharmacological inhibition of the H^+^ current leads to highly specific malformations in coccolith morphology (type-pH) that have only previously been observed in cells grown at low pH.

**Fig. 5:**
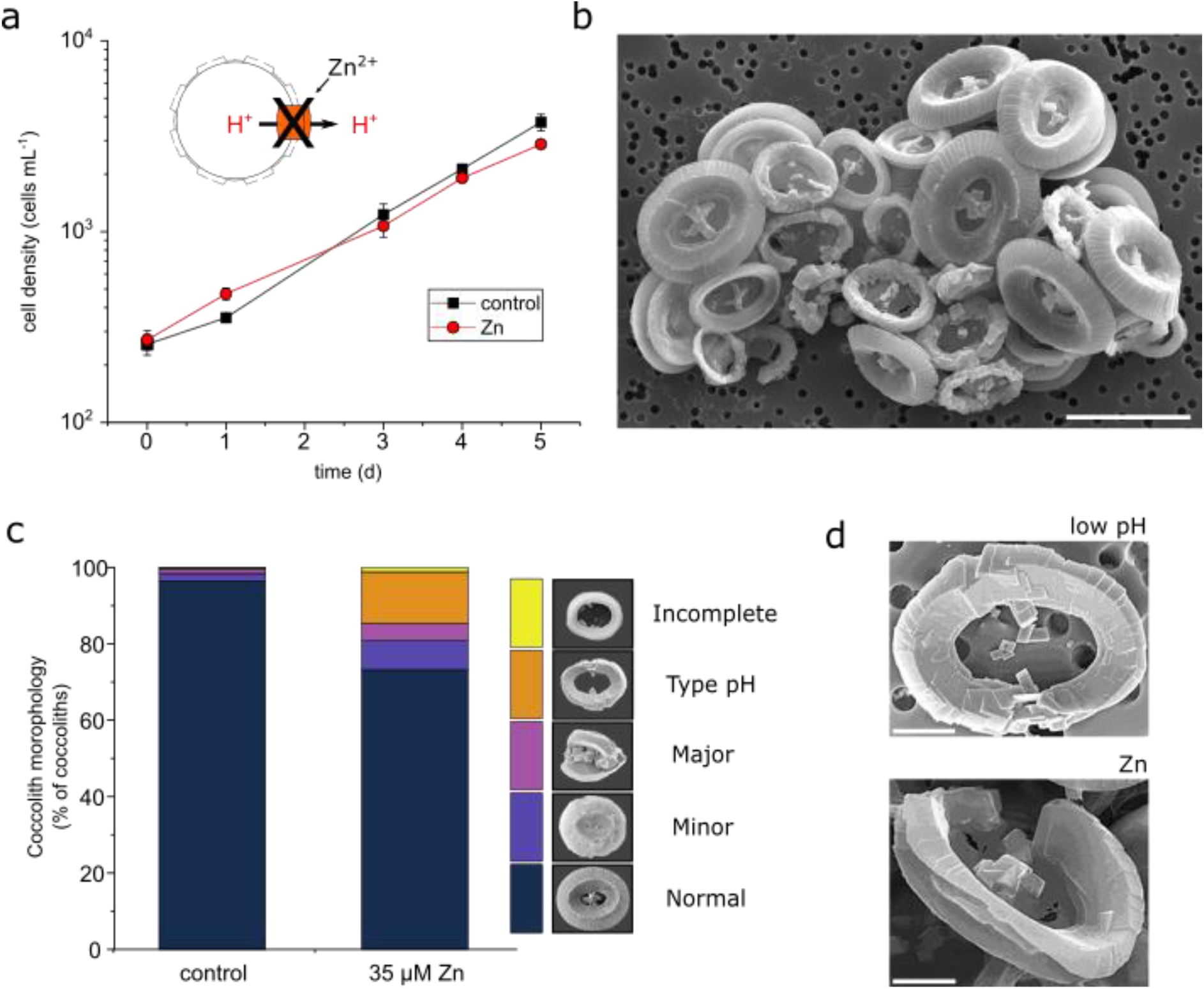
Effects of Hv inhibitors on coccolith morphology in *C. braarudii*. **a** Cell growth in the presence of the H^+^ channel inhibitor ZnCl_2_ (35 μM) at seawater pH of 8.15. n=3. Error bars = SE. Application of a similar concentration of Zn (30 μM) results in a decrease of approximately 50% in the amplitude of the outward H^+^ current ^21^. **b** SEM image of *C. braarudii* cells treated with ZnCl_2_ (35 μM) for 5 days showing the presence of many distinctive ‘type-pH’ coccolith malformations. Note also the collapse of the coccospheres due to the inability of the coccoliths to interlock. Bar = 10 μm. **c** Quantitative analysis of coccolith morphology. Coccoliths were categorised into morphological categories (see Methods). The counts represent the mean of three independent replicate treatments. A minimum of 350 coccoliths were counted for each replicate. Cells exposed to 35 μM Zn exhibit a substantial increase in the proportion of the distinctive type-pH coccolith malformations. **d** Higher resolution SEM images of type-pH morphological defects found in cells exposed to low pH (pH 7.55 from experiment described in Fig. 2) and Zn (35 μM). Bar = 2 μm.

## Discussion

We have shown that voltage-gated H^+^ channels play a critical role in pH homeostasis during coccolith formation. Hv channels are regulated by the plasma membrane H^+^ electrochemical gradient and are primed to respond to decreases in intracellular pH, allowing rapid H^+^ efflux to restore intracellular pH ^21^. However, as extracellular pH decreases to values predicted in future ocean acidification scenarios, H^+^ efflux diminishes to the extent where this mechanism is no longer effective. Under such conditions, H^+^ channel function may only be maintained by reduced pH_cyt_ or depolarisation of the membrane potential, with likely pronounced impacts on other physiological processes. Indeed, the loss of capacity to generate outward H^+^ currents shown here in cells acclimated to lower pH suggests a physiological need to switch off this mechanism of pH_cyt_ control. Our results also indicate that alternative mechanisms to maintain cytosolic pH homeostasis, such as energised forms of H^+^ transport (e.g. H^+^-ATPases, Na^+^/H^+^ exchangers), are incapable of dealing with the exceptional H^+^ load generated by intracellular calcification (Supplementary Table 2). Loss of H^+^ channel function will therefore lead to cytoplasmic acidosis unless calcification rate is reduced. The reliance on H^+^ channels for pH homeostasis may constrain the ability of coccolithophores to adapt to lower pH environments.

The inactivation of the outward H^+^ current at low seawater pH most likely involves changes in protein translation or post-translational modifications that modify channel activation. Elucidating these cellular mechanisms will be key in determining whether the loss of H^+^ channel function is reversible. Long-term experiments examining whether coccolithophores may eventually adapt to ocean acidification conditions have yielded complex results, but support a trend of decreased calcification rates ^47, 48^. Whilst physiological adaptations to low seawater pH are possible, such as recruiting alternative mechanisms for pH regulation or reducing calcification rates to lower the H^+^ load, these would either incur increased energetic costs or reduce the overall degree of calcification in future populations (Supplementary Table 2). The substantial heterogeneity in individual cell responses to low pH observed in the present study is also likely to be significant in determining how selection may operate on natural populations.

The sensitivity of *C. braarudii* calcification to low pH may not only be relevant to future ocean scenarios, but may also contribute to its distribution in current oceans. *C. braarudii* is commonly found in temperate regions, including the Iberian and Benguela upwelling systems that are associated with significant variability in seawater pH ^35, 36, 49^. It is notable that *C. braarudii* populations in the Iberian Peninsula are predominately associated with frontal systems at the periphery of the upwelled waters ^35^, rather than directly within the upwelling regions that experience the greatest extremes of seawater pH.

Calculated cellular H^+^ budgets differ considerably between coccolithophore species. In heavily-calcified species, where calcification can occur at twice the rate of photosynthesis ^10^, rapid removal of excess H^+^ is essential. However, in lightly-calcified species H^+^ production by calcification may be balanced by H^+^ consumption during photosynthesis, resulting in a much lower dependence on functional H^+^ channels. Calcification status may therefore be an important determinant in the sensitivity of coccolithophores to ocean acidification. A recent meta-analysis of multiple species revealed that the sensitivity of calcification rate to elevated seawater CO_2_ showed a strong positive correlation to PIC/POC ratio ^11^. Moreover, heavily-calcified species such as *Calcidiscus leptoporus* and *Gephyrocapsa oceanica* show highly malformed coccoliths under future ocean acidification conditions, whereas coccolith morphology in lightly-calcified species, such as *Syracosphaera pulchra, Chrysotila carterae* and *Ochrosphaera neopolitana* is less sensitive ^15, 25, 27, 47, 50^. Indeed, evidence from boron isotope approaches indicated that *O. neopolitana* is able to maintain a constant pH in the coccolith vesicle over a range of seawater carbonate chemistries, although the pH range examined was relatively narrow (pH 8.05-8.35) ^25^. Our data provide mechanistic insight into the differential sensitivity of coccolithophore species, suggesting that H^+^ load is likely to be the key determinant of their sensitivity to ocean acidification. This conclusion is seemingly at odds with observations of over-calcified morphotypes of *E. huxleyi* at higher CO_2_ levels in the Southern Ocean ^51^, and millennial-scale trends indicating a correlation between increasing prevalence of ‘over-calcified’ morphotypes of *E. huxleyi* with increased atmospheric CO_2_ over approximately the past 150 kA ^52^. However, laboratory analyses of ‘over-calcified’ *E. huxleyi* morphotypes suggest that this phenotype relates primarily to coccolith morphology rather than calcification rate, as their PIC/POC ratios are not higher than those with normal coccolith morphology ^53^.

While non-specific defects in coccolith morphology reflecting reduced calcification in response to ocean acidification have been observed in many studies ^15, 37^, the unique malformations observed here in *C. braarudii* now provide a mechanistic link between seawater pH, the ability to regulate pH_cyt_ and coccolith morphology. The highly specific nature of the *C. braarudii* malformations may facilitate the identification of low pH stress in environmental populations and aid the characterisation of past ocean acidification events in the fossil record ^37^. Modelled reconstructions indicate that, apart from the last ca 25 Ma, surface ocean pH has been lower than at present over much of the 200 Ma since the emergence of calcifying coccolithophores ^54, 55^. This suggests that coccolithophores possess some capacity to adapt to the lowering of seawater pH over geological timescales. However, the very rapid predicted decline in surface ocean pH driven by anthropogenic CO_2_ emissions may limit the degree to which coccolithophores can adapt their physiology. Recent evidence indicates that the mass extinction event at the (K-Pg) boundary (66 MYA), which led to the loss of 90% of coccolithophore species, was associated with rapid ocean acidification ^56^. It is notable that many of the coccolithophore species that survived the K-Pg Cretaceous–Paleogene mass extinction event were coastal species ^57-59^, which may have been better suited to variable seawater pH and therefore less sensitive to ocean acidification.

Multiple environmental parameters in addition to carbonate chemistry are predicted to change in future oceans, including temperature, nutrient availability and ecosystem scale changes in the abundance of predators, pathogens and competitors ^60^. Predicting the response of coccolithophore populations to future environmental change is therefore highly complex. Our incomplete understanding of the haplo-diplontic life cycle of coccolithophores further limits our ability to predict how natural populations may respond to unfavourable conditions ^61^. However, our results show that the physiology of heavily-calcified species such as *C. braarudii* is best suited to a constant seawater pH and that calcification is likely to be severely affected by ocean acidification. The ability of coccolithophores to calcify intracellularly, which has facilitated the evolution of remarkably diverse coccolith architecture, required the development of specialised physiological mechanisms for pH homeostasis that ultimately may constrain the ability of certain species to adapt to rapid changes in ocean pH.

## Methods

### Cell culturing

Cultures of *Coccolithus braarudii* (PLY182g) (formerly *Coccolithus pelagicus ssp. braarudii*) were grown in sterile-filtered seawater containing additions of nitrate, phosphate, trace metals and vitamins according to standard F/2 medium as described previously ^41^. Silicon, selenium and nickel were also supplemented in concentrations of 10 µM, 0.0025 µM and 0.0022 µM, respectively. Dilute-batch cultures were maintained at 15°C under an irradiance of 50 µmol m^- 2^ s^-1^ with a 16:8 h light:dark cycle. Cells were cultured in in autoclaved borosilicate bottles with minimal headspace and gas-tight lids to avoid in- and outgassing of CO_2_ (Duran Group, Mainz, Germany).

### Acclimation to various seawater pH

Cultures were pre-acclimated for 4 d in a range of seawater pH/carbonate chemistry conditions and then used to inoculate test cultures (Supplementary Table 1). Triplicate cultures were used for all analyses, except for the pH_cyt_ measurements where 5 replicate cultures were grown. Growth rates and coccolith morphology were determined after 5 d in test conditions (i.e. a minimum of 9 d acclimation). Physiological measurements (pH_cyt_, patch clamp) were performed between 5-10 d after inoculation into test conditions. Adjustment of seawater pH/carbonate chemistry was performed by modulating total alkalinity (TA) with amounts of HCl or NaOH at constant DIC in sealed containers. Cell density was kept between 500 and 4000 cells mL^-1^ to minimise carbonate chemistry drifts. Carbonate chemistry was measured immediately after cell inoculation and at the end of the acclimation period measuring pH_NBS_ and total alkalinity TA with a pH meter (Mettler Toledo, UK) and alkalinity titrator (TitroLine alpha plus, Schott Instruments, Germany). TA measurements were corrected with certified reference materials (CRM; provided by A. Dickson; Scripps Institution of Oceanography, U.S.A.). Data was accuracy-corrected with certified reference materials supplied by A. Dickson (Scripps Institution of Oceanography, US). Calculations were made with CO2SYS ^62^.

### Phenotypic changes in physiology

Cell growth was assessed by daily cell counting with Sedgewick Rafter counting chamber (Graticule Optics, UK) using a Leica DM 1000 LED light microscope. Specific growth rates (μ) were calculated from daily increments in cell concentrations counted every 24 or 48 h ^63^.

Cellular POC content was estimated by measuring the area of decalcified cells microscopically. The area was converted to volume, assuming cells were spherical. The mean volume [μm^3^] of at least 20 – 50 cells per culture was converted into POC quota using the equation:

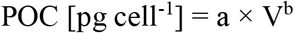

where a and b are constants (0.216 and 0.939, respectively) for non-diatom protists ^64^. The cellular PIC contents were also estimated microscopically, using the volume of the coccosphere. To obtain the cellular PIC quota, the volume of the coccolith is required. The following equation was used.

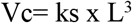

Here, Vc is the volume of coccoliths and can be estimated using coccolith length L and the shape constant ks ^65^ which is 0.06 for *C. braarudii*. The cellular PIC quota is calculated from the following equation:

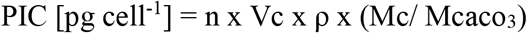

where n is the total number of coccoliths per cell including the discarded coccoliths; ρ is the calcite density of 2.7 pg µm^-3^ assuming coccoliths are pure calcite; Mc/ Mcaco_3_ is the molar mass ratio of C and CaCO_3_.

Time points for sampling cell volumes (t = 10 h after the onset of the light phase) were chosen in order present daily means (according to a modified version of the model provided in ^63^). Production rates of POC and PIC (pmol cell^-1^ d^-1^) were approximated as cellular POC content [pmol cell^-1^] × μ × [d^-1^] or cellular PIC content [pmol cell^-1^] × μ × [d^-1^], respectively. Determination of pH optima was performed by determining the vertex of a polynomial fit (second order) of three independent experiments (each experiment consists of triplicate cultures acclimated to the five different seawater pH).

### Calculation of the proton motive force

The proton motive force (pmf) at the different seawater pH was estimated as

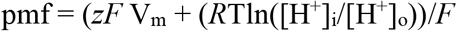

where *z* is the electrical charge of H^+^, *F* is the Faraday constant [J V^-1^ mol^-1^], V_m_ is resting membrane potential [V], *R* is the gas constant [J mol^-1^ K^-1^] and T is temperature [K]. We used a value of -46 mV for V_m_, which represents a mean of previously measured values in *C. braarudii* ^21^. To show how changes in pH_cyt_ and V_m_ may influence pmf during changes in pH_o_, we calculated pmf using two values for pH_cyt_ (6.8 and 7.1) and resting V_m_ (−46 and -28 mV).

### Calculation of H^+^ production rates

H^+^ production and consumption during photosynthesis and calcification were calculated based on POC and PIC production rates. To determine the possible range for net H^+^ load, we estimated maximal and minimal values based on a cell taking up only CO_2_ (no H^+^ consumed per C fixed during photosynthesis, 2 H^+^ generated per C precipitated during calcification) or taking up only HCO_3_^-^ (1 H^+^ consumed per C fixed during photosynthesis, 1 H^+^ generated per C precipitated during calcification).

### Scanning electron microscopy analysis of coccolith morphology

Samples for SEM analysis were filtered on polycarbonate filters (0.8 μm pore-size), dried in a drying cabinet at 50°C for 24 h, then sputter-coated with gold-palladium using an Emitech K550 sputter coater at the Plymouth Electron Microscopy Centre (PEMC). Scanning electron micrographs were produced with both Jeol JSM-6610LV and Jeol JSM-7001F at PEMC. The following categories were used to describe coccolith morphology of *C. braarudii*: normal; minor malformation (element malformation that does not impair interlocking); major malformation (shield malformation that impairs interlocking); malformation type-R (gap in both shields so that the shield elements do not form a closed oval shape); rhomb-like malformation (elements severely malformed displaying rhomb-like crystal morphology); incomplete (closed oval shape, but with incompletely grown shield elements that do not exhibit malformations); type-pH (closed oval shape, but with short shield elements exhibiting malformations) (Fig. 2). Despite a superficial similarity between the categories “incomplete” and “type-pH” that can make them difficult to distinguish in the light microscope, they are easily distinguished in SEM. An incomplete coccolith indicates that coccolith growth was stopped prematurely, but it does not indicate a malfunctioning of the morphogenetic machinery. Therefore, the label “incomplete” should only be applied to coccoliths that do not feature any malformations ^66, 67^. A malformed coccolith contains individual elements that have a disrupted morphology, rather than just an abnormal size. An average of ∼350 coccoliths was analysed per replicate culture flask, with triplicate cultures examined for each treatment ^68^. Coccolith categorization and counting employed standard methodologies as described in detail in ^66^.

### *C. braarudii* Patch-Clamp Recording and Analysis

*C. braarudii*, cells were decalcified by washing cells with Ca^2+^-free ASW containing 25 mM EGTA ^21, 41^. The recording chamber volume was 1.5 cm^3^, and solutions exchanged using gravity-fed input and suction output at a rate of 5 cm^3^ min^−1^. Patch electrodes were pulled from filamented borosilicate glass (1.5 mm OD, 0.86mm ID) using a P-97 puller (Sutter Instruments, Novato, CA, USA) to resistances of 3-6 MΩ. All external and pipette solutions are described in Supplementary Table 3. Sorbitol was added to pipette solutions to adjust the osmolarity to 1,200 mOsmol kg^−1^. Liquid junction potentials were calculated using the junction potential tool in Clampex (Molecular Devices, Sunnyvale, CA) and corrected off-line. Whole cell capacitance and seal resistance (leak) were periodically monitored during experiments by applying a <5 mV test pulse. Currents were linear leak subtracted in Clampfit (Molecular Devices, Sunnyvale, CA) using the pre-test seal resistance. Current voltage relations were determined on leak subtracted families by measuring the maximum steady state amplitude (averaging between 10 and 50 ms of the current trace). Reversal potentials were determined by manually measuring the peak tail currents of leak subtracted families of traces and calculating a linear regression versus test voltage. Series resistance was monitored throughout the experiments and whole cell currents were analysed only from recordings in which series resistance varied by less than 15%.

### Cloning of *C. braarudii HV2* into mammalian expression vector

*HV2* (CAMPEP_0183380698) from *C. braarudii* was identified by sequence similarity searches of the *C. braarudii* transcriptome (MMETSP0164) ^45^ using *C. braarudii HV1* as a query (ADM25825.1). The predicted coding sequence for *HV2* was synthesised (GenScript, Piscataway, NJ) after being codon-optimised for expression in human cells, and sub-cloned into pcDNA3.1-C-eGFP using *HindIII* and *BamHI*. A 6 bp Kozak sequence (GCCACC) was included upstream of the ATG, and the stop codon removed.

### Culturing and transfection of HEK293 Cells

HEK293 cells (ATCC CRL-1573) were cultured at 37°C in a humidified atmosphere containing 95% CO_2_ in a Dulbecco’s modified Eagle’s medium (DMEM, Gibco, 12800-017) containing 10% fetal bovine serum (Gibco™ Fetal Bovine Serum, Qualified, Cat. 26140095), 2 mM glutamine, penicillin 100 U mL^-1^ and streptomycin 100 μg mL^-1^. Cells were passaged every 3 to 4 d at 1:6 or 1:12 dilutions (cell mm^-2^). HEK293 cells were plated for transfection onto 35 mm poly-L-lysine coated glass-bottom dishes (35-mm) (www.ibidi.com). Transfections of HEK293 were performed with 1.0 µg of expression vector using Lipofectamine 2000 (ThermoFisher). After 12 to 30 h of incubation, cells were rinsed and maintained with fresh growth media until used for electrophysiological experiments. Cells exhibiting GFP fluorescence were subsequently selected for electrophysiological analysis. HEK293 cells transfected with pCDNA3.1-eGFP alone were used as a control.

### HEK293 Whole Cell Patch-Clamp Electrophysiology

Standard whole-cell patch recordings were performed at room temperature with a Multiclamp 700B amplifier (Molecular Devices, Sunnyvale, California) under the control of pClamp10 software (Molecular Devices, Sunnyvale, California). Patch electrodes were pulled from filamented borosilicate glass (1.5 mm OD, 0.86mm ID) using a P-97 puller (Sutter Instruments, Novato, CA, USA) to resistances of 3-6 MΩ. Voltage errors incurred from the liquid junction potentials (LJPs) and series resistance (recorded from the amplifier) were corrected by subtraction post hoc. These corrected voltages were used to plot IV curves and in all subsequent investigations.

### Intracellular pH Measurements

For pH_cyt_ measurements, *C. braarudii* cells were loaded with the cell-permeant acetoxymethyl ester for of the pH sensitive fluorescent dye, carboxy SNARF-1. Cells were incubated with SNARF-1 (5 μM) for 20-40 minutes, before being washed with ASW and placed in a poly-lysine coated imaging dish. Cells were imaged using a Nikon Ti Eclipse fluorescence (TIRF) system, equipped with a Photometrics Evolve EM-CCD camera and a Photometrics DV2 beamsplitter. SNARF-1 was excited between 540-560 nm and fluorescence emission was captured at 580 nm (570-590nm) and 630 nm (620-640 nm). Images were recorded at the rate of 3.3 frames s^-1^ (300 ms exposure). Background fluorescence was minimal and was therefore not subtracted.

pH_cyt_ values at acclimation conditions were derived by measuring SNARF-1 fluorescence in ASW with identical carbonate chemistry to that used for acclimation. For each treatment, pH_cyt_ was measured on a minimum of three independent days. To measure the response of cells to changes in external pH (pH_o_), acclimated cells were loaded with SNARF-1 in control ASW at pH 8.15. pH_o_ was then changed by consecutively perfusing the cells with ASW of pH 6.55, 7.55 and 8.75 for 5 minute time intervals (flow-through approximately 3 mL min^-1^ in 0.5 mL total volume). In a final step, cells were washed with ASW of pH 8.15 to determine the pH_cyt_ drift between the beginning and the end of the experiment. If the pH_cyt_ offset was > 4%, measurements were discarded from analysis.

For image processing, the mean fluorescence emission ratio (F_630_/F_580_) was determined using a region of interest encompassing the whole cell. We were unable to achieve a satisfactory *in vivo* calibration for SNARF-loaded cells using the nigericin technique, as we found that dye fluorescence was not stable after the addition of this protonophore. We therefore used an *in vitro* calibration, measuring the fluorescence emission ratio (F_630_/ F_580_) of SNARF-1 (40 µM) in buffer (130 mM KC1, l mM MgCl_2_, 15 mM HEPES) of a range of pH (pH 6.75 - 7.5). From the calibration curve, the following relation was obtained (R^2^=0.86):

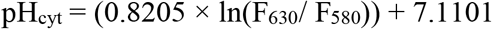

### qPCR analysis of gene expression

Quantitative reverse-transcriptase polymerase chain reaction (qPCR) was performed for *HV1* and *HV2* in cultures acclimated to pH 7.55, 8.15 and 8.75 (in triplicates). The expression of two endogenous reference genes (ERGs), *EFL* and *RPS1*, was measured alongside expression of the two target genes. Primers were designed to amplify products approximately 150 bp in length. Primer quality was tested by performing efficiency curves for serial dilutions (1:5) of each primer pair (efficiencies were > 98%, R^2^ values > 0.96). RNA was extracted from *C. braarudii* cells using the Isolate II RNA Mini Kit (Bioline) with on column DNA digestion. 30 ml of exponential growth phase culture (approximately 4,000 cells mL^-1^) was centrifuged at 3800 *g* for 5 min at 4°C. Quality and quantity of extracted RNA were tested using a Nanodrop 1000 (Thermo Fisher Scientific) (A_260_/A_280_ ratios > 1.80). cDNA was synthesised from 50 ng RNA using a SensiFAST cDNA Synthesis Kit (Bioline), with a combination of random hexamers and oligo dT primers. No Reverse-Transcriptase Controls (NRTCs) were generated to ensure no DNA contamination had occurred. qPCR runs were performed using a Rotorgene 6000 cycler (Qiagen, USA). Each reaction (20 μL) consisted of 1 μL of cDNA substrate and 19 μL of a SensiFAST No-ROX Kit Master Mix (Bioline, UK). Following primer optimisation, 0.4 μM primer were used for all genes. PCR cycles were run with Rotorgene Q series software, comprising an initial 95°C 2 min hold period, 40 cycles of 95°C denaturing for 5 s, 62°C annealing for 10 s and 72°C extension step (acquisition at end of extension step) for 20 s. A high resolution melt (HRM) curve, 72 - 95°C with 1°C ramp was conducted after amplification to ensure all amplicons had comparable melting temperatures.

For each sample, 1 μL of cDNA were analysed in technical triplicates (target genes) or duplicates (reference genes). One qPCR plate contained all *HV1* or *HV2* primer reactions (or the NRTCs), as well as the ERG reactions (*EFL* and *RPS1*), control reactions (= *HV1* expressed in the pH 8.15 acclimation), no template controls (NTC) and two positive controls. Stability of the ERG was tested using geNorm ^69^. qPCR data were analysed using a efficiency corrected DDCt method, normalizing to the geometric mean of the two ERGs ^69^. Expression of *EFL* in all NRTC was at least 10 Ct smaller than the sample.

### Phylogenetic Analyses

Previously identified Hv1 sequences from coccolithophores ^21^ were used as queries for sequence similarity searches of the available haptophyte transcriptomes within the Marine Microbial Eukaryote Sequencing Project ^45, 70^; reassembled reads NCBI accession PRJEB37117). Further Hv sequences from other representative protist lineages were obtained from the Joint Genome Institute (https://phycocosm.jgi.doe.gov/phycocosm/home). Protist Hv sequences possess an extended extracellular loop between transmembrane domains S1 and S2 ^21, 44^, enabling the generation of a longer multiple sequence alignment and improved resolution of the haptophyte Hv sequences. Hv sequences from other lineages (e.g. animals) lack the extended S1-S2 loop, although phylogenetic trees constructed with a wider range of eukaryotes exhibited a similar topology. Potential Hv sequences identified by sequence similarity searches were manually inspected using a multiple sequence alignment to assess the presence of conserved residues essential for H^+^ channel function ^23^. The multiple sequence alignments were then refined using GBLOCKS 0.91b to remove poorly aligned residues ^71^. Phylogenetic analysis was performed using the maximum likelihood method within MEGAX software ^72^ after prior determination of the best substitution model (WAG with gamma and invariant).

### Statistical analyses

For coccolith morphologies mean and SEM values were calculated from experimental replicates with a minimum of 350 coccoliths scored for each replicate. For electrophysiology means and SEM values were calculated from individual replicate cells from each treatment, with n numbers given in each Fig. For intracellular pH measurements, differences in pH between treatments were tested with 1-way ANOVA Holm-Sidak post hoc tests. Differences in distribution of pH values between treatments were assessed with 2-sample Kolmogorov-Smirnov tests.

## Data availability

All data is deposited and archived on our institutional servers or relevant genomics databases, according to institutional and funders’ requirements. Data will made available from the corresponding authors on request following publication.

## Acknowledgements

The work was supported by an ERC Advanced Grant to CB (ERC-ADG-670390) and a NERC award to GLW (NE/N011708/1). Electron microscopy analyses were performed at the PEMC (Plymouth University, UK).

## Author contributions

DK, GLW and CB conceived the study. DK was responsible for the majority of experimental analyses, with AC performing electrophysiology and GL performing the SEM analysis of coccolith morphology. KEH performed cloning of *HV2*. DK, AC and GL analysed the data. DK, GLW and CB wrote the manuscript. All data will be made available on request following publication.

The authors declare that they have no competing interests.

All data needed to evaluate the conclusions in the paper are present in the paper and/or the Supplementary Materials. The data can be provided by the corresponding authors pending scientific review and a completed material transfer agreement. Requests for data should be submitted to the corresponding authors.

