## Supplementary data for "Reduced H^+^ channel activity disrupts pH homeostasis and calcification in coccolithophores at low ocean pH"

**Supplementary Information**

**Supplementary Figures 1-8**

**Supplementary Tables**

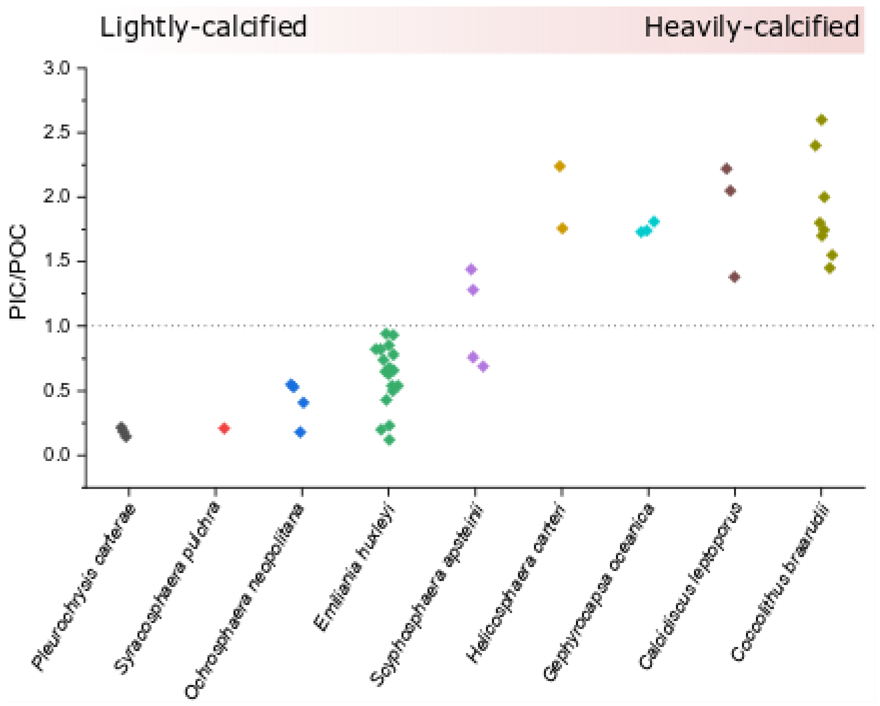


**Supplementary Figure 1. Coccolithophore species exhibit different degrees of calcification.** The graph shows the ratio of particulate inorganic carbon to particulate organic carbon measured from a range of different coccolithophore species. The data were collated from a range of studies, but do not represent an exhaustive list of all studies. In studies where carbonate chemistry was manipulated, only the PIC/POC values for cultures similar to present day carbonate chemistry scenarios have been used (pH 8.05-8.30). The data indicate that the $H^+$ load associated with intracellular calcification is likely to differ markedly between species. Note that the very low PIC/POC values in some *E. huxleyi* strains likely reflects the reduced ability to calcify exhibited by some *E. huxleyi* isolates following prolonged maintenance in laboratory culture. Data were taken from [9,11,15,21,25,27,31,33,47,74-79].

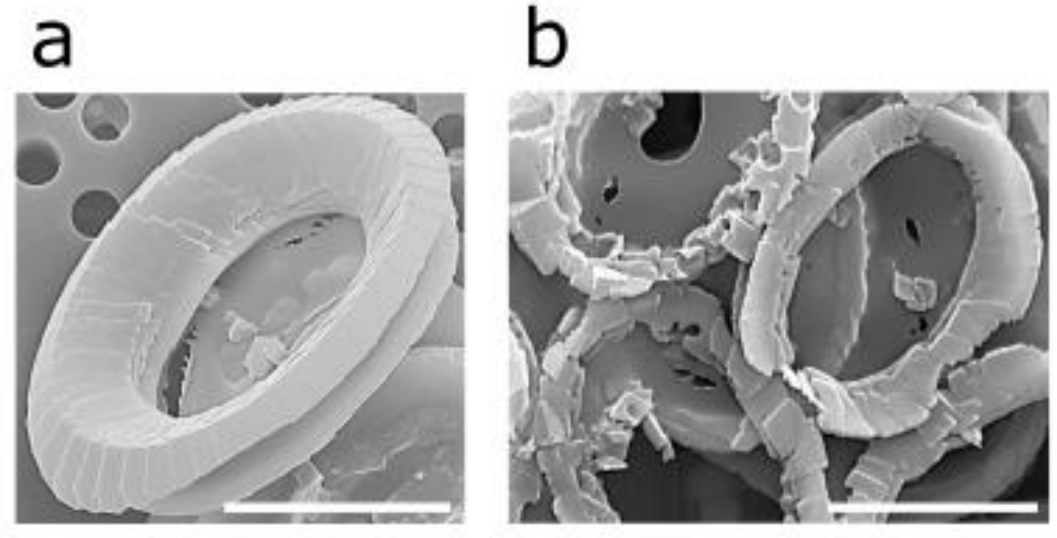


**Supplementary Figure 2. Morphological distinction between incomplete and type-pH malformations. (a)** SEM displaying incomplete coccolith. Note that the individual elements are shorter, but not malformed. Bar = 5 μm. **(b)** Type-pH malformation showing the individual elements are not only shorter, but display extensive malformations. Bar = 5 μm.

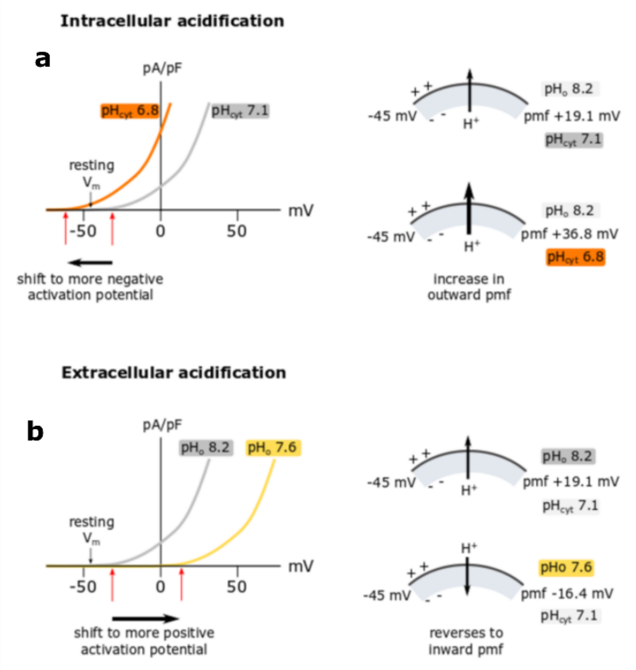


**Supplementary Figure 3. The impact of changes in the transmembrane $H^+$ gradient on the operation of voltage-gated $H^+$ channels. (a)** A small decrease in cytosolic pH ($pH_{cyt}$) due to the intracellular production of $H^+$ has a twofold effect on $H^+$ channel operation. It shifts the activation potential more negative, increasing the open probability at resting membrane potential ($V_m$) (Taylor 2011), and also increases the transmembrane $H^+$ gradient, which results in a larger outward proton motive force (pmf). These two effects combine to support $H^+$ efflux and restore $pH_{cyt}$. **(b)** If the extracellular pH is lower, as in an ocean acidification scenario, the opposite occurs. The activation potential of the $H^+$ channel is shifted to a more positive membrane potential and the pmf is reversed to a net inward pmf. In this scenario, small decreases in $pH_{cyt}$ are unlikely to activate the $H^+$ channel and the pmf is unfavourable for $H^+$ efflux. Thus, $H^+$ efflux through the $H^+$ channel is unlikely to be an effective mechanism of pH regulation.

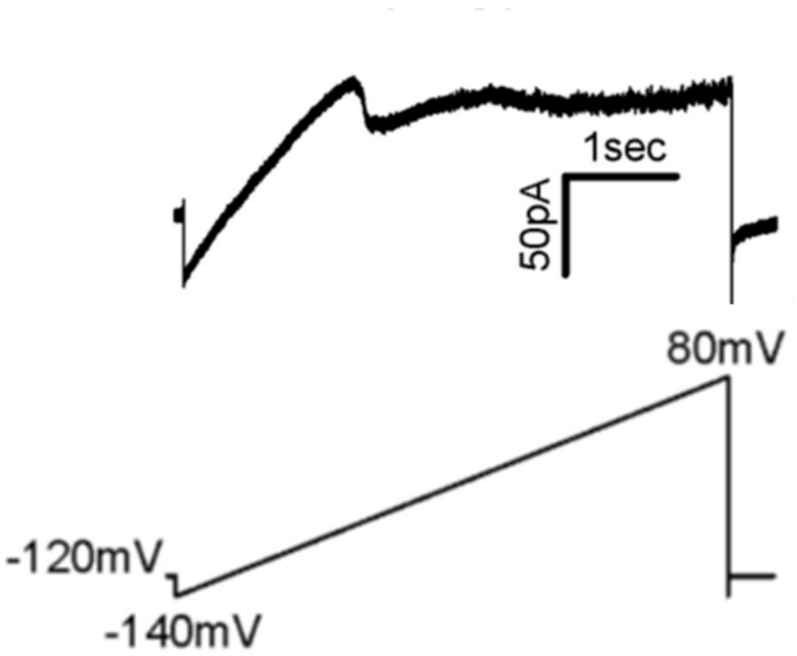


**Supplementary Figure 4. *C. braarudii* cells without an outward $H^+$ current still exhibit functional $Cl^-$ currents.**

Representative trace of a *C. braarudii* cell acclimated to pH 7.55. *Top trace:* The presence of the large inward rectifying $Cl^-$ current is clearly visible at negative membrane potentials but no outward current is apparent at positive potentials. *Bottom trace:* Using the patch-clamp technique, a voltage ramp (from − 140 mV to + 80 mV) was applied to the membrane to monitor both inward and outward current.

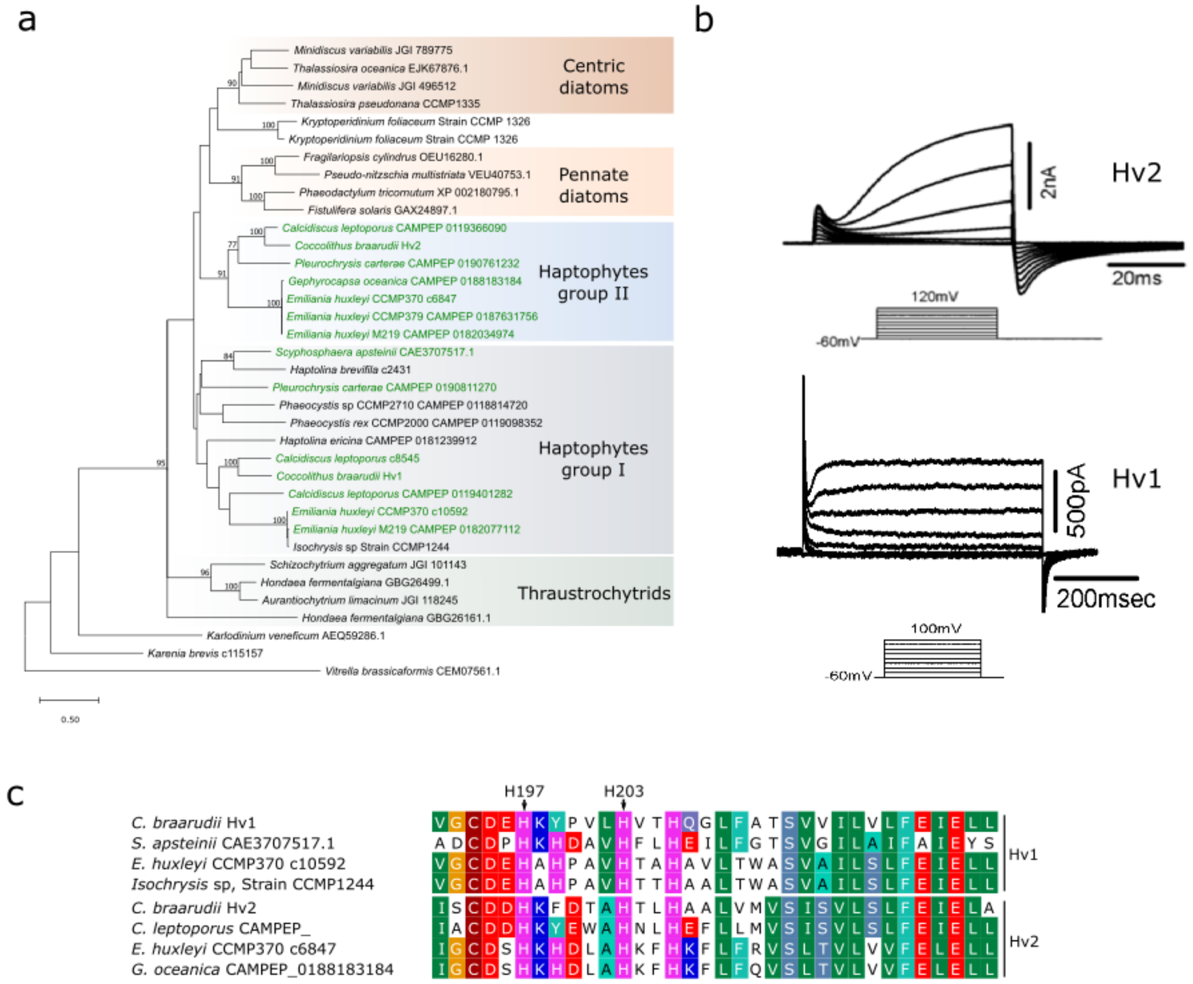


**Supplementary Figure 5. Coccolithophores possess multiple Hv homologues.** **(a)** A phylogenetic tree of haptophyte Hv homologues. A multiple sequence alignment of haptophyte Hv sequences was constructed using eukaryote Hv sequences. Protist sequences have an extended extracellular loop between transmembrane domains S-1 and S2, allowing for a longer multiple sequence alignment. Green text indicates calcified haptophyte lineages. Coccolithophore Hv sequences fall into two major groups. Group I contains Hv sequences from all haptophytes (including non-calcified haptophytes), including previously characterised Hv1 sequences from *E. huxleyi* and *C. braarudii* [21]. Group II represents a well-supported clade containing only sequences from calcified coccolithophore species. This distribution suggests that Group I sequences (Hv1) could perform a general role in haptophyte physiology (e.g. in supporting NADPH oxidase activity [42], whereas the restriction of Group II (Hv2) sequences to calcified coccolithophores suggests that they could play specialised roles in the calcification process. The tree was constructed using the maximum likelihood method with the WAG substitution model with gamma and invariant. Numbers above nodes indicate bootstrap support (100 bootstraps were performed, values>70 are shown). The final alignment size was 149 amino acids. **(b)** Heterologous characterisation of Hv2 from *C. braarudii*. To confirm that coccolithophore group II sequences also function as voltage-gated $H^+$ channels we expressed *C. braarudii* Hv2 in HEK293 cells. Patch clamp analysis revealed that *C. braarudii* Hv2 generated robust $H^+$ currents that were similar to those generated by *C. braarudii* Hv1. **(c)** Multiple sequence alignment showing that the histidine residues (arrowed) responsible for Zn inhibition of *E. huxleyi* Hv1 [21] are strongly conserved across haptophyte Hv sequences.

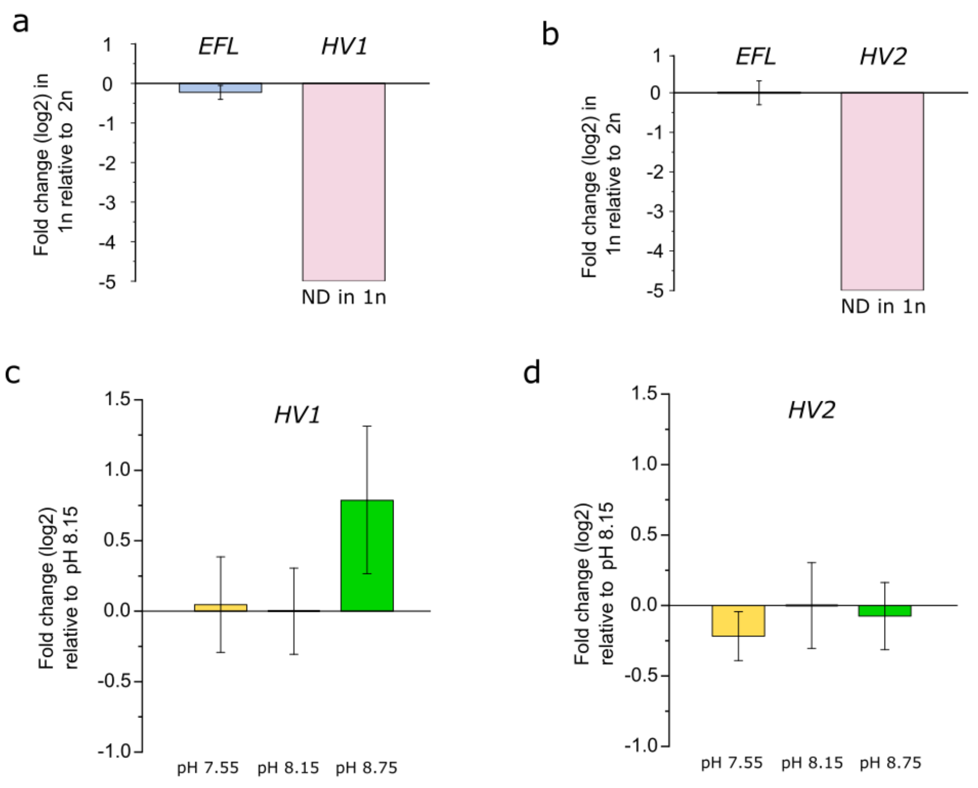


**Supplementary Figure 6. Gene expression of *C. braarudii* $H^+$ channels determined by qPCR. (a)** Expression of *HV1* in the haploid (1n) and diploid (2n) life cycle phases of *C. braarudii*. The haploid is very lightly calcified while the diploid is heavily calcified. Expression of Hv1 was normalised to a single reference gene (*RPS*) and is shown as the fold change in 1n relative to 2n. *HV1* was not detected (ND) in 1n cells. The expression of the *EFL* reference gene is also shown (normalised to *RPS*) to show that the expression of reference genes did not differ markedly between life cycle phases. **(b)** Expression of *HV2* in the haploid (1n) and diploid (2n) life cycle phases. Expression of *HV2* was normalised to a single reference gene (*RPS*) and is shown as the fold change in 1n relative to 2n. *HV2* was not detected in 1n cells. In all cases n=3 biological replicates, error bars represent SE. **(c)** Expression of *HV1* in *C. braarudii* cells acclimated to pH 7.55, 8.15 and 8.75. Expression was normalised to two reference genes (*EFL* and *RPS*) and is shown as fold change relative to the expression levels of *HV1* in pH 8.15 acclimated cells. No significant differences were found (1-way ANOVA). **(d)** Expression of *HV2*. Expression was normalised to two reference genes (*EFL* and *RPS*) and is shown as fold change relative to the expression levels of Hv1 in pH 8.15 acclimated cells. No significant differences were found (1-way ANOVA). In all cases error bars represent SE.

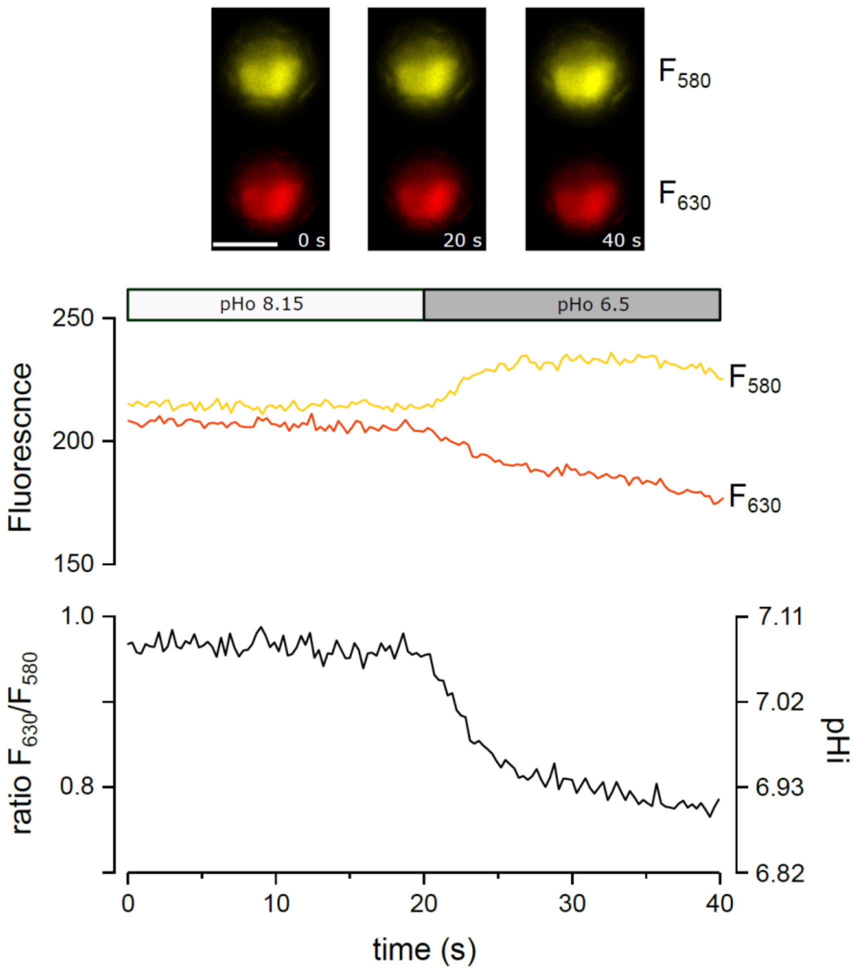


**Supplementary Figure 7. Measurement of $pH_{cyt}$ in *C. braarudii* using the fluorescent dye SNARF-1.** *C. braarudii* cells were ester-loaded with the pH-responsive fluorescent dye SNARF-1 and viewed by epifluorescent microscopy. Cells showed an even distribution of dye in the cytoplasm, indicating that the dye had not entered other cellular compartments (which could have a different pH). Fluorescence emission was monitored at 580 and 630 nm. On switching the external pH from 8.15 to 6.55 by perfusion, $F_{580}$ increases while $F_{630}$ decreases, leading to an overall decrease in the $F_{630}/F_{580}$ ratio. An *in vitro* calibration curve of SNARF-1 was used to estimate $pH_{cyt}$ (see Methods). In the example shown, the cell exhibits a rapid decrease in $pH_{cyt}$, typical of *C. braarudii* cells following exposure to lower external pH. Bar = 10 µm

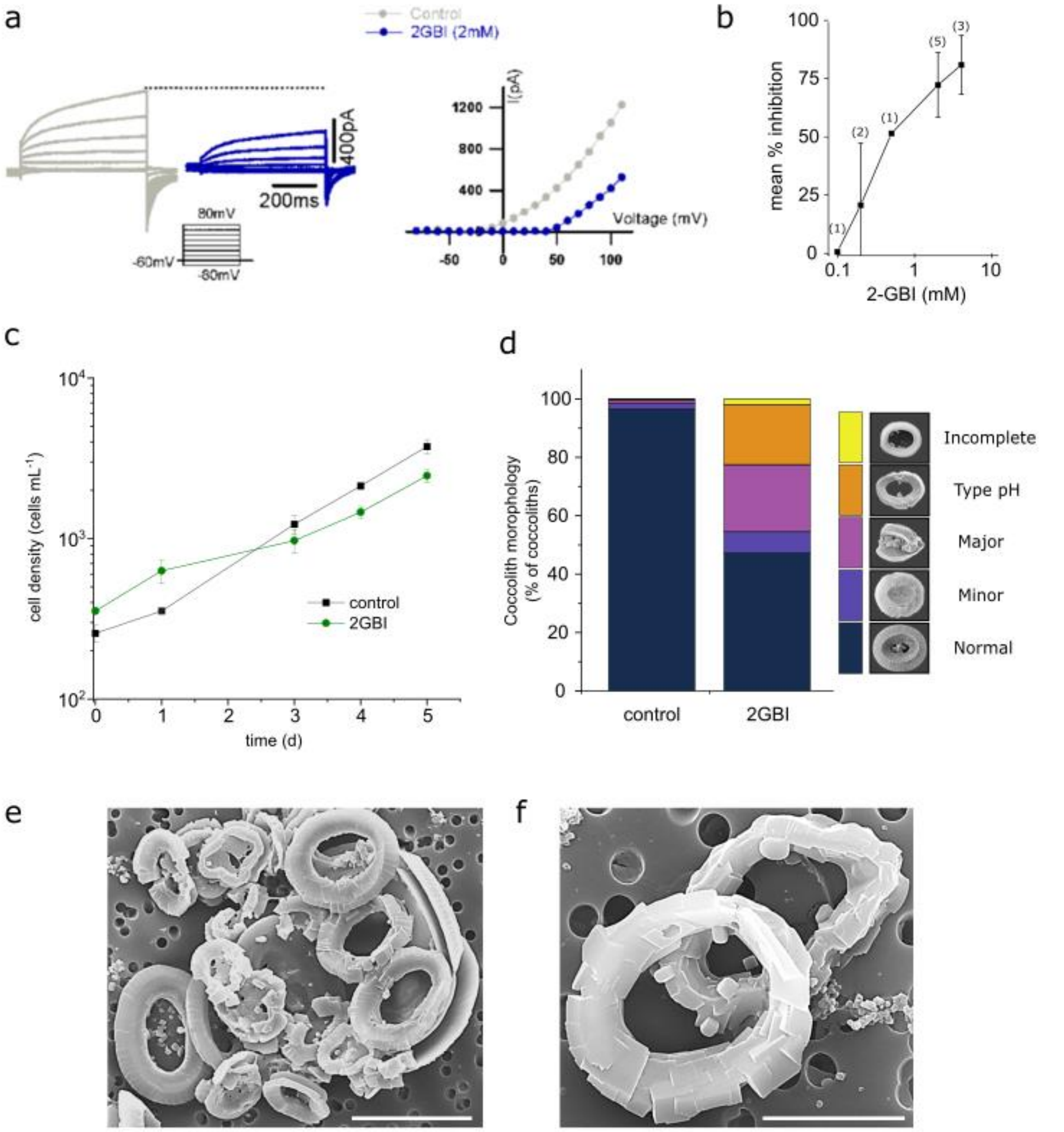


**Supplementary Figure 8. Effects of the Hv inhibitor 2-GBI on coccolith morphology in *C. braarudii* (a)** $H^+$ currents in *C. braarudii* following addition of the Hv inhibitor 2-guanidinobenzimidazole (2-GBI). 2-GBI acts by binding to the intracellular side of the voltage-sensing domain and therefore only has a limited effect on $H^+$ currents when applied externally at low concentrations in short–term experiments [46]. However, higher concentrations substantially reduced the $H^+$ current, indicating that 2-GBI is an effective inhibitor of the Hv channels in *C. braarudii*. **(b)** Outward current inhibition dose response curve of 2-GBI in *C. braarudii*. Outward currents in the presence of 0.1-4 mM 2-GBI were normalised to the untreated control currents to show percentage inhibition. Values in parentheses represent n. Error bars represent SE. **(c)** Growth of *C. braarudii* cells in 15 μM 2-GBI. Although higher concentrations of 2-GBI are required for rapid inhibition of $H^+$ currents, much lower concentrations had an effect on growth. We interpret this as a gradual internalisation of 2-GBI, which has a low membrane permeability. Cell growth is shown in the presence of 2-GBI (15 μM) at seawater pH of 8.15. n=3, error bars = SE. **(d)** Quantitative analysis of coccolith morphology following treatment with 2-GBI. Coccoliths were categorised into morphological categories (see Materials and Methods). The counts represent the mean of three independent replicate treatments, with a minimum of 350 coccoliths were counted for each replicate. Cells exposed to 15 μM 2-GBI for 5 d exhibit a substantial increase in the proportion of the distinctive type-pH coccolith malformations, observed at low pH or after treatment with the Hv inhibitor ZnCl. **(e)** SEM image of a *C. braarudii* cell treated with 2-GBI (15 μM) for 5 days showing the presence of many type-pH coccolith malformations. The inability of the coccoliths to interlock leads to the collapse of the coccosphere during preparation for SEM imaging. Bar = 10 μm. **(f)** Higher magnification SEM image of type-pH malformations induced by 15 μM 2-GBI. Bar = 5 μm.

### Supplementary Tables

#### Supplementary Table 1. Acclimation carbonate chemistry

| Accl. pH | | $pH_{NBS}$ | $H^+$ | $pCO_2$ | $CO_2$ | $HCO_3^-$ | $CO_3^{2-}$ | DIC | TA | Ω |
|---|---|---|---|---|---|---|---|---|---|---|
| 7.55 | $t_0$ | 7.56 ± 0.02 | 37 ± 1 | 1608 ± 63 | 60 ± 2 | 1920 ± 11 | 40 ± 1 | 2020 ± 12 | 2058 ± 11 | 1.0 ± 0.0 |
| | $t_{fin}$ | 7.63 ± 0.05 | 32 ± 3 | 1405 ± 174 | 52 ± 6 | 1931 ± 72 | 47 ± 2 | 2031 ± 77 | 2055 ± 67 | 1.1 ± 0.1 |
| 7.85 | $t_0$ | 7.80 ± 0.02 | 21 ± 1 | 957 ± 51 | 36 ± 3 | 1987 ± 11 | 72 ± 2 | 2095 ± 12 | 2205 ± 12 | 1.8 ± 0.1 |
| | $t_{fin}$ | 7.82 | 26 | 1231 | 46 | 2097 | 63 | 2206 | 2259 | 1.5 |
| 8.15 | $t_0$ | 8.14 ± 0.01 | 10 ± 0 | 423 ± 9 | 16 ± 1 | 1928 ± 11 | 153 ± 5 | 2097 ± 16 | 2346 ± 23 | 3.6 ± 0.1 |
| | $t_{fin}$ | 8.01 ± 0.03 | 13 ± 1 | 562 ± 41 | 21 ± 3 | 1882 ± 1 | 111 ± 9 | 2014 ± 6 | 2166 ± 20 | 2.6 ± 0.2 |
| 8.45 | $t_0$ | 8.43 ± 0.01 | 5 ± 0 | 205 ± 2 | 8 ± 0 | 1825 ± 11 | 283 ± 6 | 2115 ± 17 | 2556 ± 24 | 6.7 ± 0.1 |
| | $t_{fin}$ | 8.27 ± 0.03 | 7 ± 0 | 300 ± 21 | 11 ± 1 | 1819 ± 25 | 194 ± 12 | 2024 ± 25 | 2306 ± 34 | 4.6 ± 0.3 |
| 8.75 | $t_0$ | 8.75 ± 0.01 | 2 ± 0 | 87 ± 1 | 3 ± 0 | 1596 ± 9 | 511 ± 10 | 2110 ± 19 | 2863 ± 31 | 12.2 ± 0.2 |
| | $t_{fin}$ | 8.60 ± 0.01 | 3 ± 0 | 136 ± 2 | 5 ± 0 | 1781 ± 11 | 407 ± 11 | 2192 ± 20 | 2761 ± 33 | 9.7 ± 0.3 |

n=3 in all cases except pH 7.85 $t_{fin}$ where n = 1 due to samples lost during analysis.

**Supplementary Table 2. Potential adaptive responses of *C. braarudii* following loss of $H^+$ channel function**

| Potential adaptation | Possible consequences |
|---|---|
| **1) Restore $H^+$ channel function** | |
| Reduce $pH_{cyt}$ | Maintaining a lower $pH_{cyt}$ will help to restore $H^+$ electrochemical gradient, but is likely to interfere with many aspects of metabolism |
| Increase Vm | Depolarisation of the resting $V_m$ will help to restore the $H^+$ electrochemical gradient, but will have major consequences for many other aspects of membrane transport. $H^+$ efflux through $H^+$ channels may be feasible through repetitive transient depolarisations of $V_m$ (action potentials), although slow activation kinetics of Hv channels are not ideally suited to rapid activation. |
| Modify $H^+$ channel gating | Adaptive shift of the activation potential to a more negative $V_m$ will allow $H^+$ channels to operate at lower external pH. However, $H^+$ channel activity remains constrained by pmf, as outward $H^+$ electrochemical gradient is still required for $H^+$ efflux. |
| **2) Reduce $H^+$ load** | |
| Uptake of $HCO_3^-$ rather than $CO_2$ | Greater $HCO_3^-$ uptake will result in increased consumption of $H^+$, but $HCO_3^-$ transport is energetically more costly than $CO_2$ uptake. Cells would not benefit from elevated seawater $CO_2$. |
| Lower calcification rate | A lower calcification rate will reduce intracellular $H^+$ production, but may prevent formation of a complete coccosphere, or interfere with other aspects of coccolith function (e.g. reduce ability of *C. braarudii* placoliths to interlock) that are critical for ecological success. |
| **3) Use alternative mechanisms for $H^+$ efflux** | |
| Use energised $H^+$ transport | Use of energised mechanisms of $H^+$ transport (e.g. $Na^+/H^+$ exchange or $H^+$-ATPase) will allow $H^+$ efflux against an unfavourable $H^+$ electrochemical gradient, although energised $H^+$ transport is unlikely to match the capacity for rapid $H^+$ efflux that is required in heavily calcified species and would have increased energetic costs. |

**Supplementary Table 3. External and pipette solution compositions for patch clamp recording of *C. braaudii* and HEK293 cells.**

| | *C. braarudii* | | HEK293 cells | |
|---|---|---|---|---|
| | External solution (mM) | Internal solution (mM) | External solution (mM) | Internal solution (mM) |
| **NaCl** | 450 | | 140 | 10 |
| **KCl** | 8 | | 5 | 115 |
| **$CaCl_2$** | 10 | | 2 | 0.1 |
| **$MgCl_2$** | 30 | 5 | 1 | 10 |
| **$MgSO_4$** | 16 | | | |
| **$NaHCO_3$** | 2 | | | |
| **HEPES** | 20 | 100 | 10 | 20 |
| **K-glutamate** | | 200 | | |
| **EGTA** | | 5 | | 1 |
| **pH** | 8.15 | 7.5 | 7.4 | 7.2 |
